# Horse, not zebra: accounting for lineage abundance in maximum likelihood phylogenetics

**DOI:** 10.64898/2026.03.25.714173

**Authors:** Nicola De Maio

## Abstract

Maximum likelihood phylogenetic methods are popular approaches for estimating evolutionary histories from genome data. These methods do not make prior assumptions regarding strategies used for deciding which genomes were sequenced. However, in genomic epidemiology the sequencing rate is often agnostic to the specific pathogen strain considered. In this scenario, a pathogen strain prevalence should be reflected in its relative abundance in the genome data. Here, I show that this simple assumption, when appropriate and incorporated within maximum likelihood phylogenetics, greatly improves the accuracy of phylogenetic inference.

I introduce and assess two separate approaches to achieve this. The first approach rescales the likelihood of a phylogenetic tree by the number of distinct binary topologies obtainable by arbitrarily resolving multifurcations in the tree. This approach interprets multifurcations as the result of lack of signal for resolving a bifurcating topology rather than as an instantaneous multifurcating event. The second approach instead includes a tree prior that assumes that genomes are sequenced at a rate proportional to their abundance.

Both approaches favor phylogenetic placement at abundant lineages, and dramatically improve the accuracy of phylogenetic inference in scenarios like SARS-CoV-2 phylogenetics, where large multifurcations are common. This considerable impact is also observed in real pandemic-scale SARS-CoV-2 genome data, where accounting for lineage prevalence reduces phylogenetic uncertainty by around one order of magnitude. Both approaches were implemented in the open source phylogenetic software MAPLE v0.7.5.4 (https://github.com/NicolaDM/MAPLE).

## 1 Introduction

Molecular phylogenetic methods are fundamental to genomic epidemiology, and are used for many applications beyond basic reconstruction of phylogenetic trees, including the inference of the origin of outbreaks[1, 2], transmission histories[3–5], geographical spread[6–8], prevalence[9–11], variant fitness[12–14], and lineage assignment[15, 16], to name a few. Maximum likelihood phylogenetic methods[17–19] are generally accurate, versatile, and practical, and make no prior assumption regarding the shape of the tree and the sampling/sequencing process. These methods have in fact usually been developed for general analyses in evolutionary biology, where sequences are included based on their availability, and where the abundance of sequences within a lineage is not necessarily representative of the richness of the lineage itself, but might be affected by other factors, such as the ease with which species within the lineage are recognized, collected and sequenced, and the perceived usefulness of these sequences. In genomic epidemiology, instead, not only the same exact pathogen genome might be sampled multiple times from different hosts, but often we can assume that the abundance of a pathogen strain at a certain time and location is reflected in the number of genomes of that strain sequenced at the given time and location (Fig. 1). This feature of genomic epidemiological datasets is typically not used in phylogenetic analyses, but could it be used to improve phylogenetic inference accuracy?

**Figure 1:**
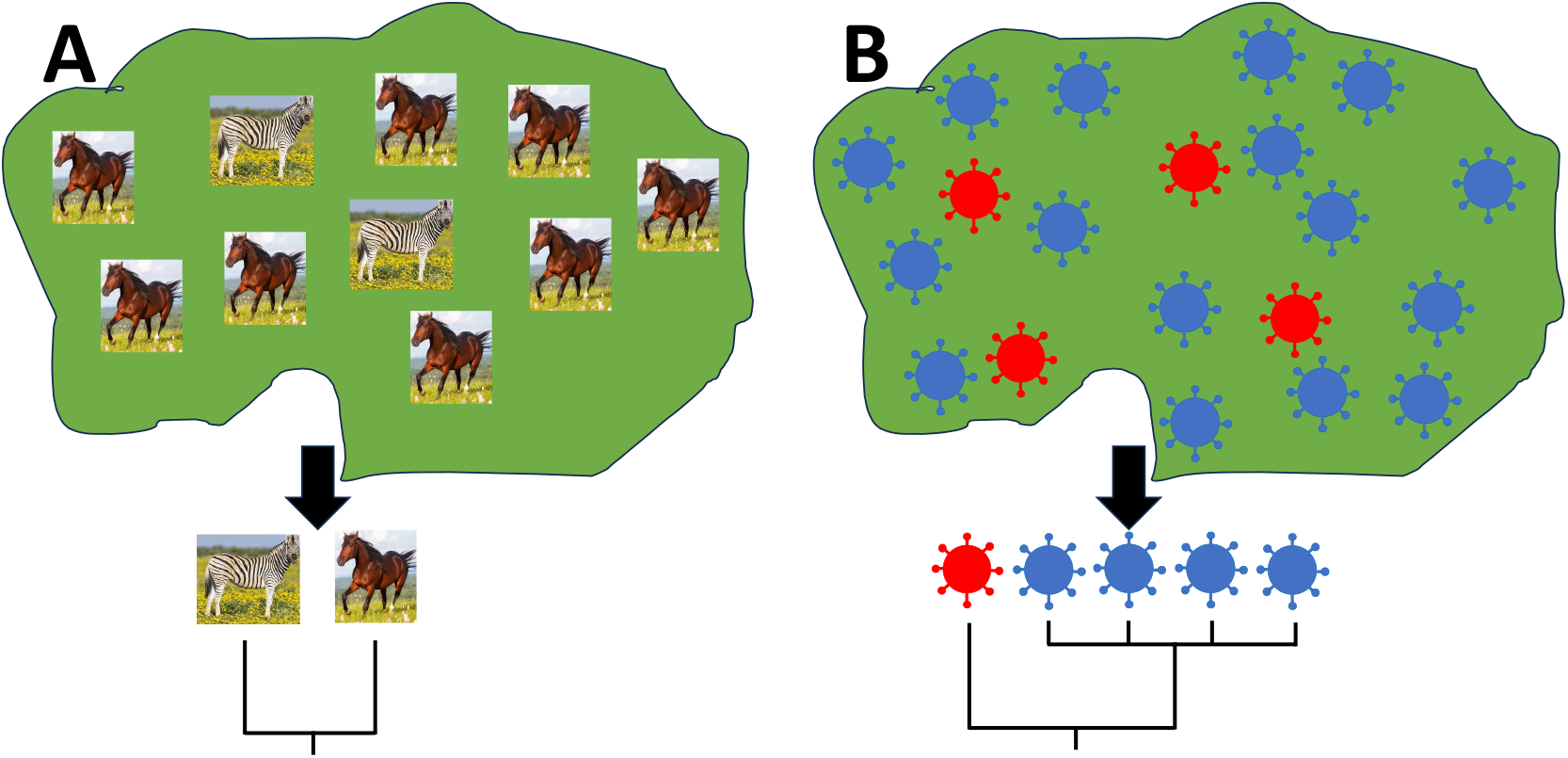
Graphical representation of the hypothesis of prevalence-driven genome abundance. **A)** In evolutionary biology, one generally cannot assume that the abundance of individuals in a species, or of species within a lineage, is reflected in the proportion of sequences from that species/lineage included in the analyses. In the graphic, horses are four times more abundant than zebras, but each species is represented by only one genome in this example sampling. **B)** In genomic epidemiology, the strain of the considered pathogen often does not impact the probability of sampling, and so it is usually expected that the prevalence of a strain is reflected in the abundance of sequenced genomes from that strain. In the graphic the blue strain is four times more abundant than the red strain, and consequently four times more blue strain genomes are sampled and sequenced than red strain genomes. Example phylogenetic trees relating the samples are shown under each sub-figure. In the medical community, rare diseases are often referred to as “zebras”, and common ones as “horses”, and a common principle is that “when you hear hoofbeats, think of horses, not zebras”, meaning that all symptoms being equal, the diagnosis of a common disease should be prioritized over the diagnosis of a rare disease[20].

As a simple example, consider two complete genomes *G*_1_ and *G*_2_ differing at only one genome position *i*, and consider an incomplete genome *G*_3_ (e.g. due to amplicon dropout[21]) that does not contain genetic information at position *i* but that is otherwise identical to both *G*_1_ and *G*_2_. Both phylogenetic placements of *G*_3_, at *G*_1_ or *G*_2_, result in the same likelihood, and so they cannot be distinguished by maximum likelihood (see Fig. 2). However, if genome *G*_1_ is more abundant than *G*_2_ in the population where samples are collected from, and if we assume that the abundance of genomes in this population is reflected in the abundance of the genomes in the considered dataset and phylogeny, then we have to conclude that *G*_3_ more probably represents an additional copy of *G*_1_ than a copy of *G*_2_.

**Figure 2:**
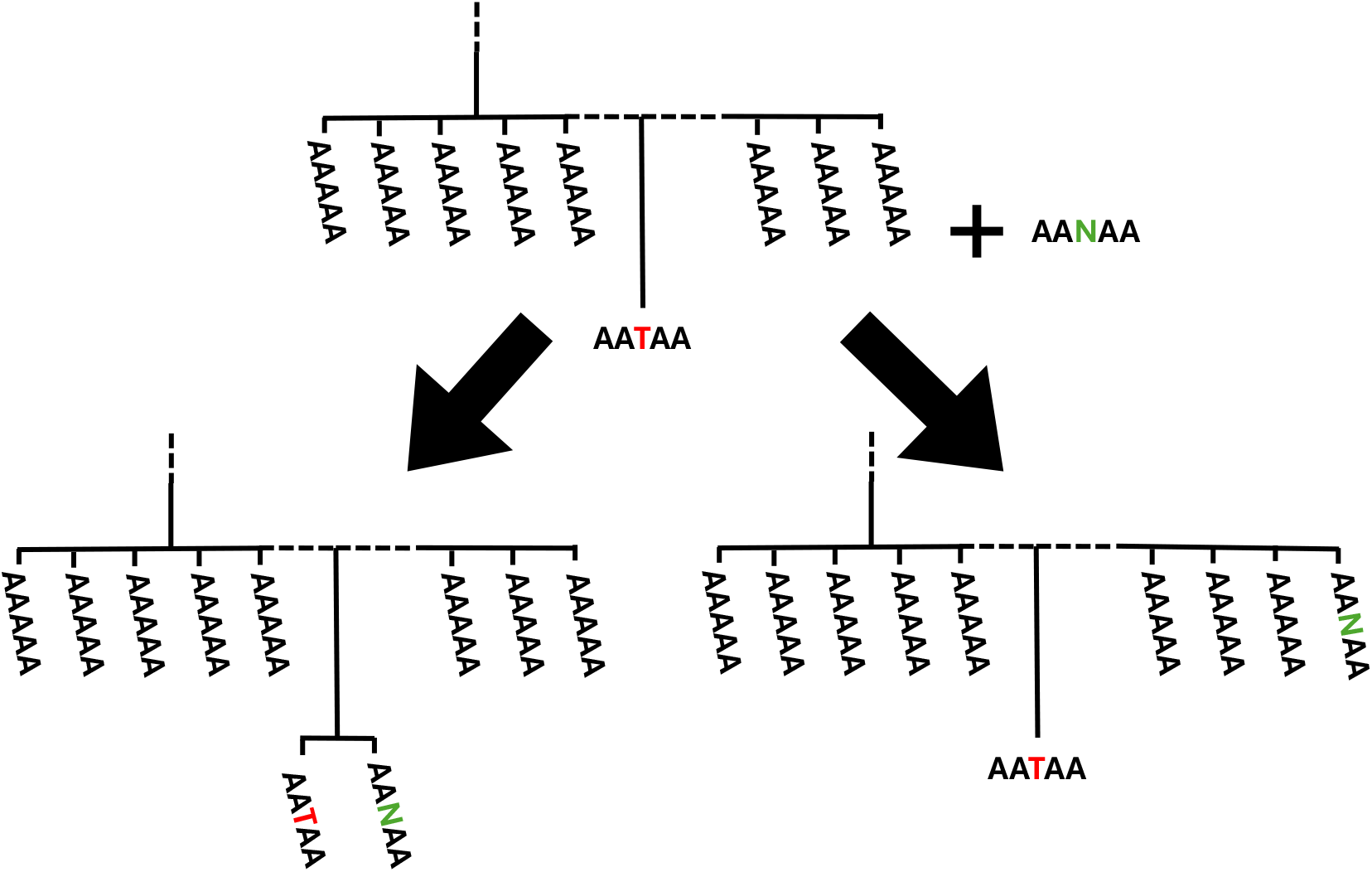
Example of resolvable phylogenetic uncertainty. In the top part of the plot I show part of an example phylogenetic tree. For conciseness, I use dashed lines to represent parts of the tree that are not drawn, and I consider example genomes of length 5bp. Genome *G*_1_=AAAAA is represented in the tree by many samples, leading to a large multifurcation. Its descendant genome *G*_2_=AATAA has only been sampled once. Let us consider the problem of placing the incomplete sequence *G*_3_=AANAA onto the tree. Here N represents a non-informative genome position. Placing *G*_3_ with *G*_2_, like in the bottom left subplot, or with *G*_1_, like in the bottom right subplot, results in the same phylogenetic likelihood, so these two placements are not distinguishable in classical maximum likelihood phylogenetics. However, since *G*_1_ has been sampled more often than *G*_2_, and assuming that genome abundance in the data is representative of abundance in the sampled population, one should conclude that *G*_3_ is more probable to be a further sample of *G*_1_ than *G*_2_.

This idea is similar to the famous medical principle “when you hear hoofbeats, think of horses, not zebras” popularized by Theodore Woodward, who practiced in an area where horses were more common than zebras; the meaning of this principle is that, all other evidence being equal, a common disease diagnosis (a “horse”) should be considered more probable than a rare disease diagnosis (a “zebra”)[20]. A partly similar principle has been implemented in the pandemic-scale phylogenetic placement method UShER[22], which, between placements with equal parsimony scores, favors placement at a node that is the root of a larger subtree.

I will focus on the application of this principle to phylogenetic inference in genomic epidemiology, and in particular consider the case of pandemic-scale SARS-CoV-2 genome data analysis, since the high sampling density and low evolutionary distances cause many replicate genomes to be sampled and many phylogenetic tree multifurcations to be inferred[23]. Furthermore, the elevated uncertainty[24] and complexity[23] seen in SARS-CoV-2 phylogenetic inference mean that extra information in the form of lineage abundance could substantially increase the accuracy of phylogenetic tree reconstruction.

I will introduce two separate approaches inspired by these ideas. The first one, HnZ1, is based on the principle that a phylogenetic multifurcation represents a number of possible transmission histories, and so the placement of a sample at a multifurcation actually represents a number of possible alternative placements within the transmission or timed tree. The second approach, HnZ2, is based on the principle that a multifurcation in a phylogenetic tree represents a prevalent genome, and this genome has a higher a priori probability of being sampled again than a less prevalent one. I show with simulations and real data analyses that these methods improve the accuracy and reduce the uncertainty in pandemic-scale phylogenetic inference.

## 2 Results

### 2.1 New approaches

I propose two approaches to account for lineage abundance in maximum likelihood phylogenetics. I refer to them collectively as “HnZ” (“Horse not Zebra”) and individually as respectively “HnZ1” and “HnZ2”. Both approaches introduce a multiplicative factor for the phylogenetic likelihood similar to a tree prior within a maximum a posteriori inference framework, and do not affect the calculation of the phylogenetic likelihood. These components favor the placement of subtrees as descendants of large multifurcations during subtree prune and regraft (SPR) search in maximum likelihood phylogenetic inference[25], and therefore lead to inferred trees with larger multifurcations. Both approaches are implemented in MAPLE v0.7.5.4 (see Section 4.1 for more details of the implementation) available from https://github.com/NicolaDM/MAPLE.

#### 2.1.1 HnZ1

Lack of mutations on branches of a (true or simulated) phylogenetic timed tree or epidemiological transmission tree cause 0-length inferred phylogenetic tree branches, and cause uncertainty in the inferred tree topology. This uncertainty and 0-length branches are usually represented as multifurcations in the inferred tree by maximum likelihood phylogenetic methods. In the following, I will call these “mutational multifurcations” (MMs) since they represent multifurcations in the inferred mutation-annotated tree, but do not necessarily correspond to multifurcations in a timed tree or transmission tree. One can of course interpret MMs trees literally, that is, as instantaneous events in which a multiple lineage splits occur. Instead, here I interpret an MM as a class of multiple possible bifurcating topologies consistent with the considered multifurcation (Fig. 3).

**Figure 3:**
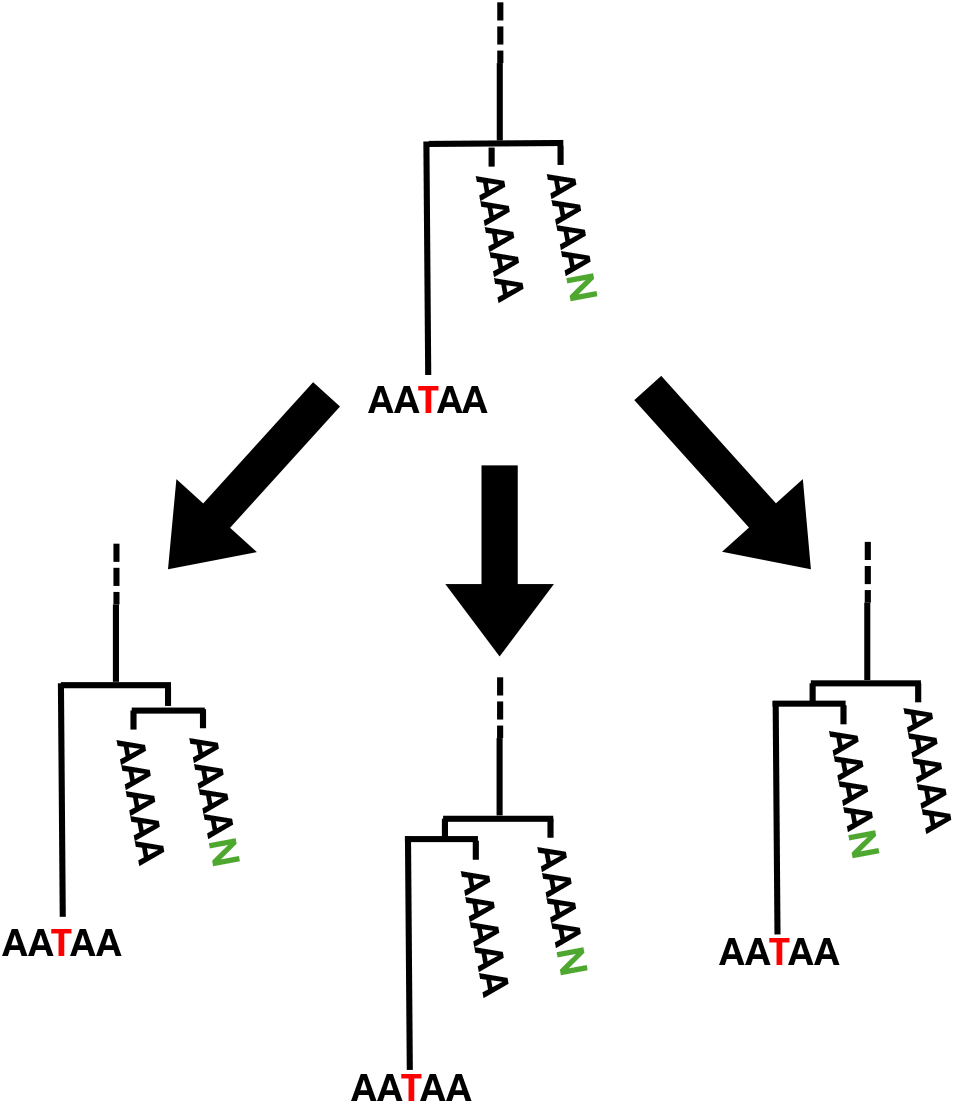
Example of bifurcating resolutions of a multifurcation. In the top is shown part of an example phylogenetic tree, similarly to Fig. 2. This subtree relates three samples, all descending from the same internal node, a mutational multifurcation (MM) of size *n* = 3. There are *H*(3) = 3 bifurcating resolutions (see Eq. 1) of this MM, all represented in the bottom part of the Figure. The short branches in the bifurcating resolutions are shown of positive length to aid visualization, but would typically have length 0 unless interpreted as branches of a timed tree.

A justification of this interpretation can be given when considering a typical Bayesian phylogenetic inference method. These methods, in the absence of mutations on a branch, do not infer a branch of length 0, but consider a distribution of short but positive branch lengths (i.e. times) consistent with the observation of 0 mutations on the branch. Similarly, in the presence of an MM, a Bayesian approach typically evaluates the bifurcating resolutions of the MM, weighing them according to their prior probability and likelihood. When considering the uncertain placement of a sample onto the phylogenetic tree, a Bayesian approach would similarly consider multiple placements within the branches resolving an MM as separate placements, and as such, unlike maximum likelihood approaches, it would tipically favor sample placement at large MMs: the total probability mass of Bayesian placement within a large MM is the sum of the probabilities of the many different individual placements within resolutions of the MM.

This is the inspiration for HnZ1: as a simplification, let us assume that different bifurcating topologies have similar priors, and that all bifurcating resolutions of an MM have similar likelihood. The phylogenetic likelihood is therefore scaled, within maximum likelihood phylogenetic inference, by the number of bifurcating topologies consistent with the given tree, so that the rescaled likelihood now represents the cumulative probability mass of all these resolutions.

The number of bifurcating resolutions of a mutationally-multifurcating tree is the product of the number of possible bifurcating resolutions for each MM in the tree. Consider a node of size *n* ≥ 1, which is defined as the number of branches in the multifurcating mutational tree descending from it (including possibly any 0-length branches connecting the node directly to samples). This node is a multifurcation if *n >* 2, and a terminal or mid-branch node if *n* = 1 (assuming the convention that terminal nodes have size 1). The number of bifurcating resolutions for a node of size *n* is the same as the number of rooted trees with *n* tips[26]:

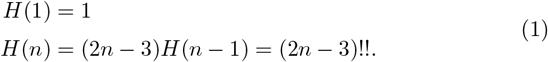

The HnZ1 component of a given multifurcating tree is then defined as the product of *H*(*n*) over all nodes in the tree (see e.g. Fig. 4).

**Figure 4:**
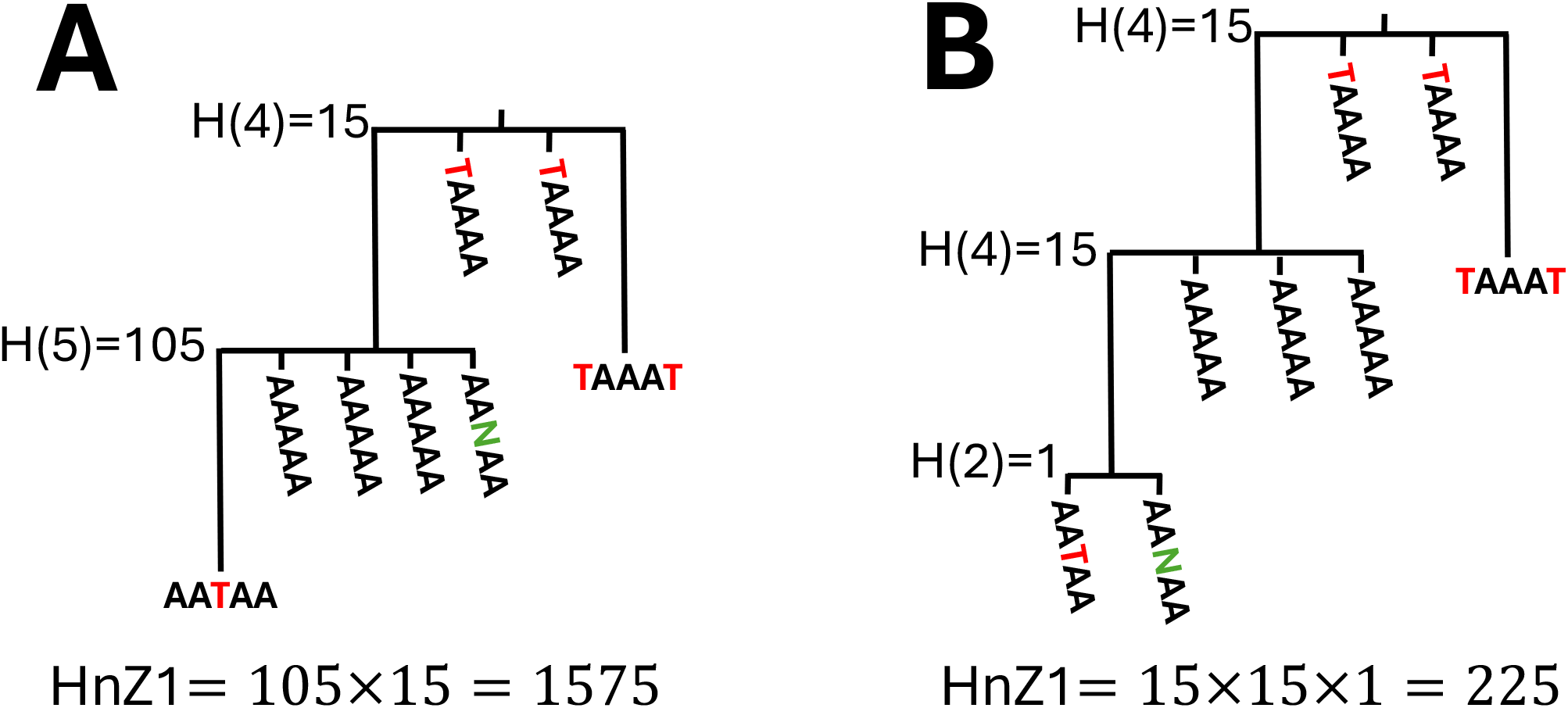
Examples of HNZ1 score calculation. **A** The HnZ1 score of this phylogenetic tree is given by the product of *H*(*n*) (from Eq. 1) over the two internal nodes in the tree (“dummy” nodes resolving MMs, which is those of 0 length not connected directly to a sample, have to be ignored), the root one of size 4 (corresponding to genome TAAAA) and the other of size 5 (corresponding to genome AAAAA). **B** Tree obtained re-placing sequence AANAA at the location of genome AATAA. This reduces the size of the MM of genome AAAAA and causes the HnZ1 score of the tree to become 7 times smaller. This shows how HnZ1 favors placement at larger MMs.

To investigate the potential impact of HnZ1 on phylogenetic inference, let us consider the problem of adding one sample onto an existing multifurcating phylogenetic tree. If the sample is placed as a direct descendant of a node of post-placement size *n*, the HnZ1 score of the tree increases by a factor of *H*(*n*)*/H*(*n* − 1) = 2*n* − 3, and therefore HnZ1 favors placements onto larger multifurcations (see Fig. 5).

**Figure 5:**
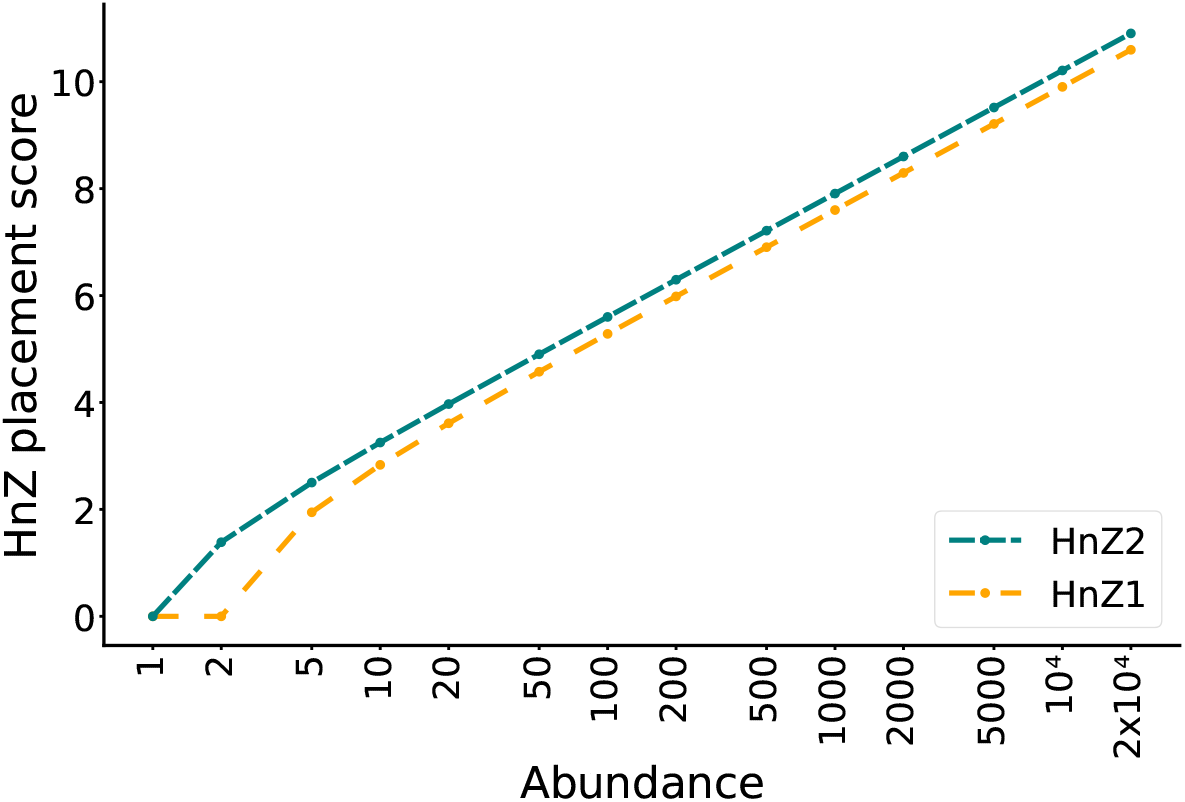
Effect of sample placement on HnZ scores. On the Y axis is shown *log*(*H*(*n*)*/H*(*n* − 1)) where *H* is one of the HnZ scores (Eqs. 1 and 2) as a function of node size *n*. Node size is defined as the number of non-dummy branches (those of positive length and those of 0 length connecting an internal node directly to a sample) descending directly from a node (internal branches of length 0 are not considered part of the tree). A multifurcation is a node of size *n >* 2. The Y axis value *log*(*H*(*n*)*/H*(*n* − 1)) represents the increase in HnZ log-score following the placement of a sample as a direct descendant of a node of size *n* − 1 before the placement. The increasing value of *log*(*H*(*n*)*/H*(*n* − 1)) as a function of *n* means that HnZ1 and HnZ2 favor the placement of samples onto larger multifurcations.

When evaluating an SPR move during phylogenetic inference, changes in *H*(*n*) are similarly calculated for all the nodes whose size is affected by the topology change (that is, either by the pruning of the considered subtree or by its regrafting) to calculate the change in HnZ1 score caused by the SPR move. To decide if to accept an SPR move (and similarly for a sample placement) the total score of the tree before the SPR move is compared to the total score of the tree after the SPR move. This total score is the product of the HNZ1 tree score and the classical phylogenetic tree likelihood score. If the total score after the SPR move is higher than the one before the SPR move, then the SPR move is accepted.

#### 2.1.2 HnZ2

The second approach, HnZ2, is also inspired from Bayesian phylogenetics. In Bayesian phylogenetics, tree priors can help distinguish between trees with similar likelihood. For example, a tree prior modeling different sampling rates across locations or times (e.g. [11, 27]) would favor sample placement at locations or times with high sampling rate and prevalence. Inspired by this, I define a simple tree prior that uses the size of mutational multifurcations to define the abundance of genomes in the tree. This measure of abundance conflates two different aspects: on one hand the number of times that a genome (corresponding to the considered node) has been sampled (corresponding to the number of samples separated by a branch of length 0 from the considered node), and on the other hand the number of times that a genome is the direct ancestor of another genome in the tree (corresponding to the number of non-0 length branches descending directly from the node). The firs aspect, the number of times a genome is sampled, is affected both by the prevalence of that genome (the number of hosts infected by that strain) and by the sampling rate at the locations and times in which the genome was prevalent. The second aspect, the number of non-dummy branches descending from a node (those of positive mutational length and those of 0 mutational length connecting an internal node directly to a sample), is also affected by the genome prevalence, but less affected by the sampling rate; for example, an early highly prevalent genome might not have been sampled at all, but might still give rise to a large multifurcation if sampling rates became higher later on in the considered outbreak. Node size *n* conflates these two aspects, and merges both the propensity of a genome to have been sampled, and the propensity of the genome of being the direct ancestor of other genomes represented by nodes in the tree. Both aspects are important for phylogenetic inference, since the first can be used to guide the placement of samples, and the second the placement of subtrees during SPR tree search.

The HnZ2 tree prior is defined as 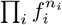 where the product is over each node *i* in the considered tree, *f*_*i*_ is the abundance of the genome corresponding to node *i*, and *n*_*i*_ is the size of node *i*. This prior represents the sampling probability (both in terms of sample collection and ancestor choice) of the genomes in the tree. Genome abundance is also approximated using the node size, and defined as *f*_*i*_ = *n*_*i*_*/N* with *N* the total number of genomes considered (accounting for multiplicity). In practice, however, the factor 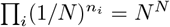 does not depend on the particular tree considered, and so can be ignored; the HnZ2 score of a node of size *n* can therefore be defined as

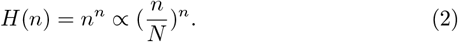

Taking the product of *H*(*n*) across all nodes in the tree gives us the total HnZ2 tree score, which is proportional to the aforementioned approximate tree prior. This total HnZ2 score is then multiplied to the classical phylogenetic tree likelihood, transforming maximum likelihood phylogenetic inference basically into maximum a posteriori inference.

Similarly to HnZ1, HnZ2 also favor sample and subtree placements at large multifurcations, but it is slightly more “aggressive” than HnZ1 (Fig. 5), in the sense that HnZ2 provides a slightly greater incentive than HnZ1 for inferring larger multifurcations.

### 2.2 Simulation-based benchmark

Here I assess the performance of HnZ1 and HnZ2 using large-scale simulated SARS-CoV-2 genome data (see Section 4.3.2 for details of the simulations). The study of SARS-CoV-2 evolution is a particularly interesting application of these methods since the dense sampling of the virus causes many large multifurcations[23], and since genome sequencing incompleteness and mutation rate heterogeneity cause elevated phylogenetic uncertainty[24, 28, 29], which HnZ1 and HnZ2 might help resolve.

#### 2.2.1 Accuracy

HnZ1 and HnZ2 greatly improve the accuracy of phylogenetic inference (Fig. 6), with HnZ1 leading to slightly higher accuracy than HnZ2, and preventing around 40% of topological inference errors compared to not using HnZ1 or HnZ2.

**Figure 6:**
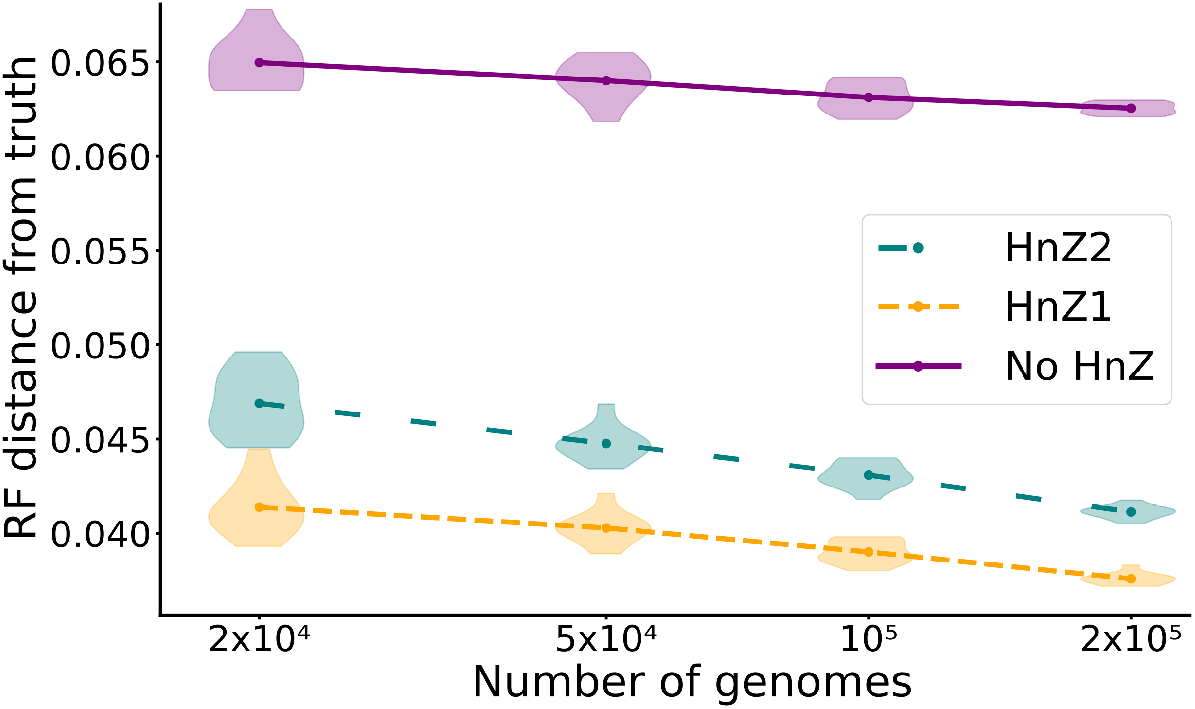
Accuracy of phylogenetic tree inference from simulated SARS-CoV-2 genome data. The Y axis shows the Robinson-Foulds distances between inferred trees and simulated trees: lower values represent more accurate phylogenetic reconstructions. On the X axis is the number of simulated SARS-CoV-2 genomes (see Section 4.3.2) included in each replicate. For each dataset size considered 10 replicates were run. Dots show the means across replicates, while violin plots show variation between replicates. For all three methods compared MAPLE v0.7.5.4 was used with an UNREST substitution model and with rate variation[29].

#### 2.2.2 Computational demand

HnZ1 and HnZ2 require additional calculations to track the size of tree nodes and to calculate HnZ scores. These calculations can be performed efficiently (section 4.1), but the usage of HnZ1 and HnZ2 still increases the time demand of phylogenetic inference, almost doubling it (Fig. 7A). One reason is that, with HnZ, less informative genomes cannot be removed from the analysis (see [30]) since different placements of these genomes, even when they have 0 phylogenetic log-likelihood cost, can still differ in HnZ score, and can affect node sizes, and therefore phylogenetic inference. Another reason seems to be that phylogenetic inference with HnZ scores requires longer SPR searches to reach convergence. HnZ scores lead instead to only small increases in memory demand (Fig. 7B).

**Figure 7:**
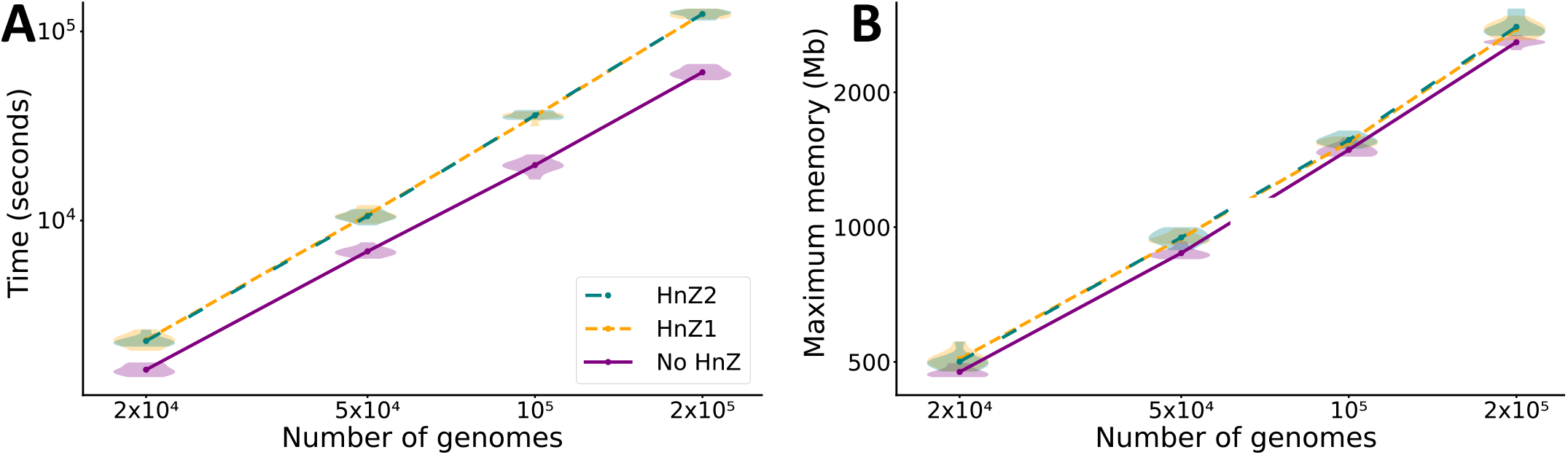
Computational demand of phylogenetic tree inference from simulated SARS-CoV-2 genome data. **A** Time and **B** maximum RAM usage of phylogenetic tree inference with MAPLE 0.7.5.4 with and without HnZ1 and HnZ2. All analyses were run on one core of an Intel Xeon Gold 6252 processor @ 2.10 GHz. All other details are as in Fig. 6.

### 2.3 Inference of SARS-CoV-2 evolution

A global SARS-CoV-2 phylogenetic tree from 2,072,111 public genomes (see Section 4.3.1) was inferred using HnZ1. Inference of an initial tree required 8.5 days and 20GB of memory with a single core, while topological improvement search parallelized over 14 cores of an Intel Xeon Gold 6252 processor @ 2.10 GHz, took around 15.5 days and 500GB. Post hoc SPRTA[24] mutational support assessment (which measures the statistical confidence in the inferred evolutionary history) with a single core required around 12 days and 30GB. Here I present the results of the comparison of the resulting tree with that inferred without HnZ (but using the same substitution model and tree search) in [24].

HnZ1 dramatically reduces phylogenetic uncertainty. Of 1,864,187 substitutions inferred with HnZ1, only ≈ 1.04% (19,356) have support *<* 50%, compared to ≈ 6.91% without HnZ. The reason is likely that HnZ1 can assign very different probabilities to otherwise equally parsimonious (and therefore similarly probable) phylogenetic placements (e.g. Fig. 2). This reduction in uncertainty affects terminal tree branches more than internal branches. With HnZ1, of 424,005 internal tree branches only ≈ 0.68% (2,921) have support *<* 50%, compared to ≈ 6.53% without HnZ. On the other hand, HnZ1 reduces the proportion of highly uncertain (support *<* 50%) positive-length terminal branches from ≈ 8.39% to ≈ 0.11%, and of 0-length terminal branches from ≈ 8.03% to ≈ 0.05%. The increased support of branches with positive length translates to a reduced uncertainty in the virus evolutionary history. The dramatic increase in support for 0-length terminal branches represents a massive reduction in uncertainty of the placement of samples that, due to sequence data incompleteness, are possibly identical to more than one other genome in the dataset (see Fig. 2 for a simplified example).

HnZ1 also appears to increase the total number of inferred mutations from 1,827,786 to 1,864,187 suggesting cases were less parsimonious placements at larger size nodes are favored by HnZ1 over more parsimonious placements at smaller size nodes.

The evolution of the AY.4 Delta sub-lineage (one of the most abundant SARS-CoV-2 lineages, represented here by *>* 480, 000 genomes) was previously described as very phylogenetically uncertain[24]. Mutation T17040C appeared ancestral to most AY.4 genomes, but frequent downstream C17040T reversions and further downstream T17040C re-reversions appeared to make phylogenetic inference very uncertain[24]. (Here a T17040C mutation is called a “re-reversion” if it occurs downstream of a C17040T reversion.) In my new analyses presented here, HnZ1 substantially impacts the inferred evolution of this lineage, leading to the inference of only 40 C17040T (from 655 without HnZ) and 41 T17040C (from 120 without HnZ) substitutions across the whole tree (Fig. 8). The evolutionary history of AY.4 inferred with HnZ1 is also much simpler, and while it still contains C17040T reversions, it does not contain any T17040C re-reversions. The uncertainty in this inference is also greatly reduced since, while without HnZ major AY.4 sub-clades have SPRTA support *<* 10%, with HnZ all major AY.4 sub-clades have 100% support.

**Figure 8:**
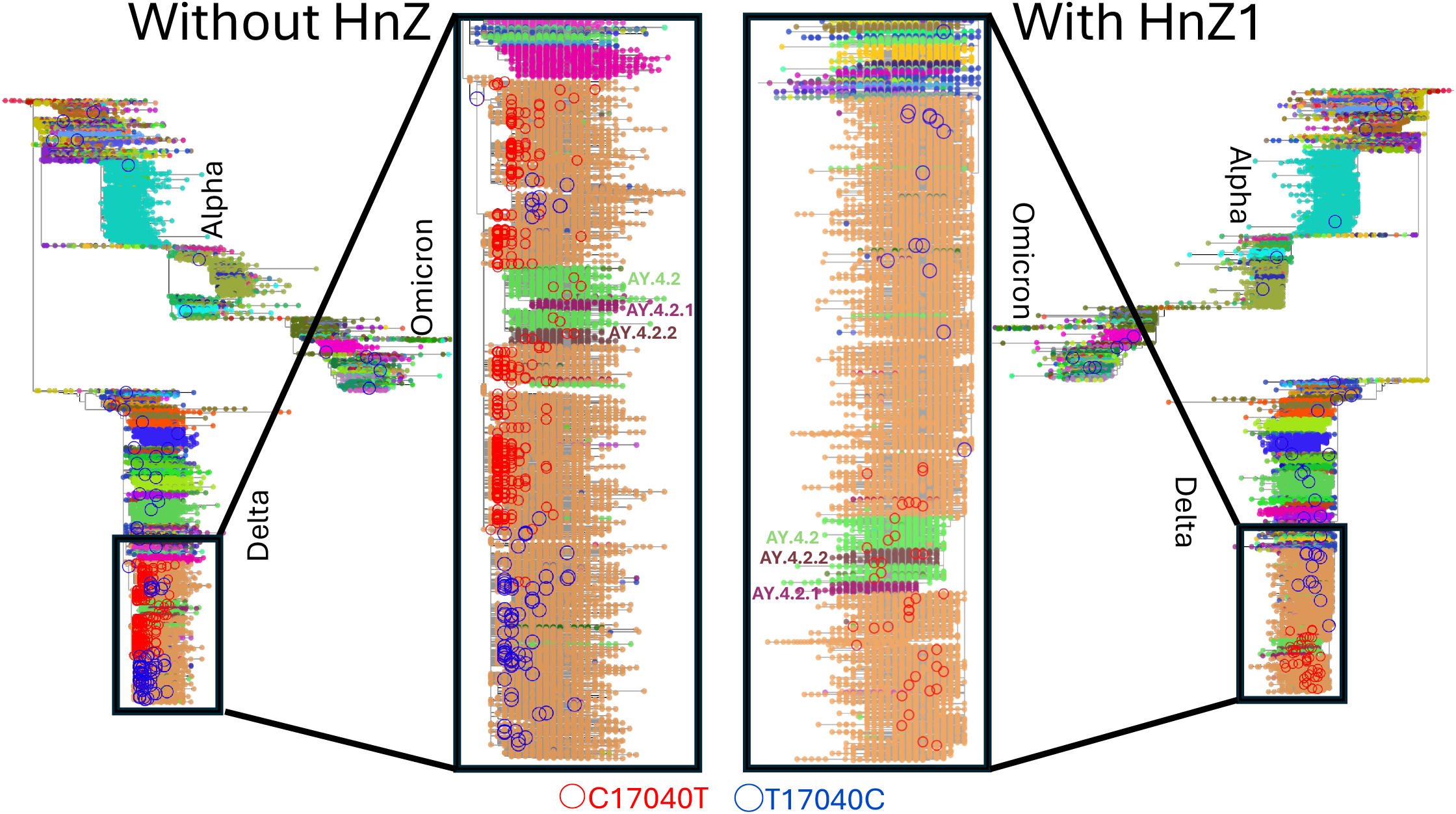
Differences between the evolutionary histories of SARS-CoV-2 lineage AY.4 inferred with and without HnZ1. On the left is the global SARS-CoV-2 phylogenetic tree inferred by MAPLE without HnZ and visualized in Taxonium[31]. Phylogenetic tips are colored according to the Pango[15] lineage assigned by Pangolin[32] v4.3 (with pangolin-data v1.21) to the corresponding genomes. The names of some lineages are also shown. The 655 inferred C17040T substitutions are highlighted with red circles, while the 120 inferred T17040C substitutions are highlighted with blue circles. The zoom-in focuses on the evolution of the AY.4 lineage. On the right is the same phylogeny (and zoom-in) inferred with HnZ1, which contains only 40 C17040T substitution and 41 T17040 substitutions. See Fig. 9 for an explanation of these differences.

The evolutionary history inferred without HnZ is hard to justify: while the many C17040T reversion could be justified by an elevated C17040T mutation rate or fitness, the high recurrence of T17040C re-reversion cannot be explained as easily, since T17040C substitutions are not as frequent outside of the AY.4 lineage. These patterns can instead be easily explained assuming that the phylogeny inferred with HnZ1 is correct, as exemplified in Fig. 9. Assuming in fact that there are two very prevalent genomes separated only by 1 substitution (in this case T17040C as on the right of Fig. 8), then many mutations are expected to occur in both genomic backgrounds, and often the same mutation will occur in both background, as in Fig. 9A. Maximum likelihood inference then struggles to distinguish between possible evolutionary histories as in Fig. 9B,C, causing inference errors, uncertainty, and reversions. HnZ will instead favor the inference of the Fig. 9A tree because it is the only maximally parsimonious one where all mutations occur in a high-prevalence genomic background, leading to less uncertainty, fewer reversions and presumably fewer errors.

**Figure 9:**
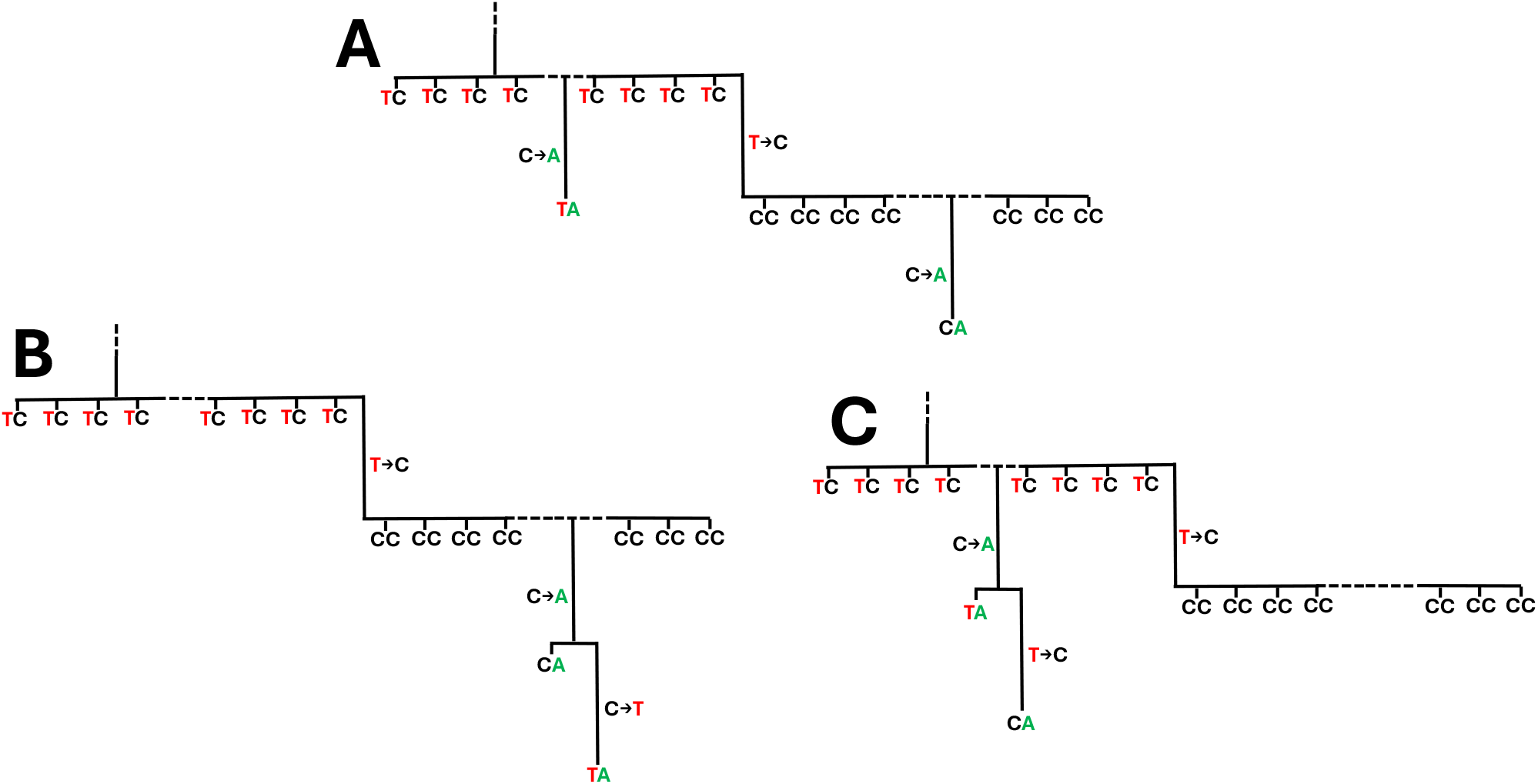
Example scenario in which HnZ impacts evolutionary inference. Here I focus on the inference of the evolutionary history of 4 genomes, which for simplicity are represented as sequences of length 2 containing only the 2 SNPs of relevance to this inference. An abundant **TC** genome (its abundance is reflected in the many samples in the phylogeny containing its sequence) is assumed to be ancestral to the other three thanks to information in the rest of the tree (not shown here, and represented by the dashed line at the top of the trees in the figure). The other three genomes are an abundant **CC** and two rare (represented only by one sample each) **TA** and **CA** genomes. Genomes **CC** and **TC** might be present in thousands of samples, and may be ancestral to many other lineages - here for graphical simplicity I use horizontal dashed lines to skip over such parts of the tree. **A)** The tree that would typically be inferred with HnZ in this scenario. Here each mutation occurs in prevalent genomic background, giving higher HnZ score. This is typically more realistic, because, for example, it accounts for the fact that the mutation **C** → **A** might have occurred at any of the many hosts infected with the **CC** genome. This example represents a typical scenario in SARS-COV-2: the **T** → **C** substitution represents the T17040C inferred to be ancestral to a large proportion of AY.4 genomes in the right side zoom-in of Fig. 8. **B)** Tree that might be inferred in this scenario by maximum likelihood without HnZ; in this case a **C** → **T** reversion is inferred instead of a second **C** → **A** mutation. This **C** → **T** reversion (representing many C17040T reversions in the left side zoom-in of Fig. 8) is inferred to have occurred in a rare genomic background (**CA**). This history might be favored by non-HnZ maximum likelihood inference if for example the **C** → **T** substitution rate is high, which is the case for SARS-CoV-2[28], which helps explain the large number of C17040T reversions in the left side zoom-in of Fig. 8. **C)** Another tree that might be inferred in this scenario by maximum likelihood without HnZ; in this case a second **T**→**C** substitution in a rare genomic background is inferred instead of a second **C**→**A** substitution in a prevalent genomic background. This history might be favored by non-HnZ maximum likelihood inference if for example the **T**→**C** substitution rate is high.

The differences in the inferred evolutionary histories of site 17040 substantially impact the inferred substitution rate at the site, which is 3.6 times higher than average when using HnZ1, and 31.9 times higher when not using HnZ. Site 17040 is in fact the one with the largest difference in inferred rate with and without HnZ. The next one is 21595, with inferred rates 3.4 times higher than average when using HnZ1, and 13.6 times higher when not using HnZ. Investigating the discrepancies between the two phylogenies shows a similar pattern to 17040: in the HnZ1 tree there are two predominant genomes within Omicron lineage BA.1.1 separated by a C21595T substitution, and while the evolutionary history of this lineage inferred with HnZ1 is relatively simple, the one inferred without HnZ shows many reversions and many additional C21595T and T21595C substitutions (Fig. 10). Another similar pattern is also observed at site 8302 (inferred rates 1.8 vs. 8.8), which appears related to a substantial BA.1.1 sub-lineage defined by an A8302G substitution. Only another 9 sites have an inferred rate difference larger than 2, suggesting that, while present in other lineages, these patterns do not typically have such a large impact as those related to T17040C in AY.4 and C21595T in BA.1.1.

**Figure 10:**
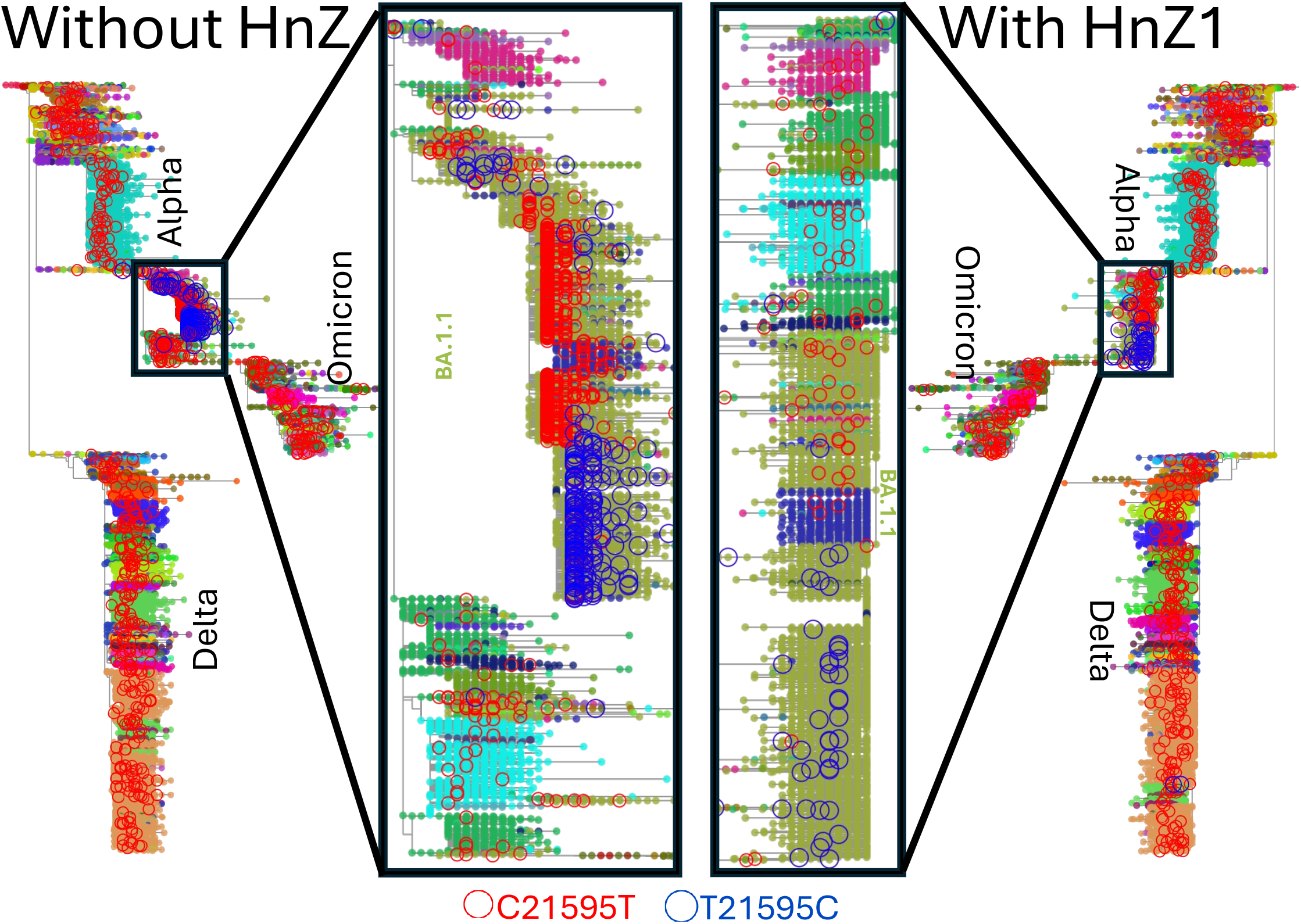
Differences between the evolutionary histories of SARS-CoV-2 lineage BA.1.1 inferred with and without HnZ1. SARS-CoV-2 phylogenies and distributions of C21595T (red circles) and T21595C (blue circles) substitutions inferred without HnZ (left) and with HnZ1 (right). The zoom-ins highlight the inferred evolution of lineage BA.1.1. On the left there are 2,003 C21595T and 255 T21595C substitutions, while on the right there are 529 C21595T and 47 T21595C substitutions. All other details are as in Fig. 8.

## 3 Discussion

Maximum likelihood phylogenetics typically does not use information regarding lineage abundance to improve tree or placement inference. Bayesian approaches, on the other hand, can implicitly account for lineage abundance by considering many similar placements, by considering bifurcating resolution of mutational multifurcations, and by using tree priors. Bayesian phylogenetic approaches are however typically substantially more computationally demanding than maximum likelihood ones, and do not concisely summarize multiple possible similar placements and topologies as single trees with mutational multifurcations.

Here I proposed two alternative approaches, HnZ1 and HnZ2, to make use of these principles and “Bayesianize” maximum likelihood inference. Using SARS-CoV-2 phylogenetics as an example, I have shown that accounting for lineage abundance can greatly increase the accuracy of low-divergence phylogenetic inference at limited additional computational demand. Compared to full Bayesian inference, for example using Monte Carlo Markov Chain (MCMC)[33], these HnZ approaches are not only more scalable, but also offer a more interpretable representation of phylogenetic uncertainty by collapsing mutational multifurcations whose bifurcating resolution is extremely uncertain, and by representing mutational uncertainty with extra probabilistically-weighed edges in the phylogenetic tree using SPRTA[24].

I have shown that HnZ substantially reduces phylogenetic uncertainty as measured using the SPRTA framework[24]. Similar reductions are expected in other measures of phylogenetic uncertainty. While the two proposed HnZ approaches can be used in the assessment of phylogenetic uncertainty, they do not however themselves account for phylogenetic uncertainty, in the sense that HnZ node scores are based on node sizes and that these sizes only account for the considered phylogenetic structure and not for its uncertainty. It would be interesting in the future to investigate approaches to alleviate this limitation while avoiding the computational costs of largescale phylogenetic MCMC inference.

HnZ appraches are expected to have a significant impact not only broadly in genomic epidemiology, but also in other scenarios with dense sampling, including when the same sequence can be represented by multiple samples, and assuming that genomes are sampled approximately proportionally to their abundance. Some applications might include for example phylogenetic placement of sequencing reads in metagenomics[34, 35], single-cell genomics[36] and cancer genomics [37]. In the future, by allowing for accuracy and scalability, these HnZ approaches have also the potential to greatly improve the accuracy of many downstream phylogenetic analyses, from lineage assignment[32], to large scale phylodynamics and phylogeography with maximum likelihood (e.g. [38, 39]) and machine learning (e.g. [40, 41]).

## 4 Methods

### 4.1 Implementation of HnZ

HnZ1 and HnZ2 were implemented within MAPLE v0.6.12 and later versions (https://github.com/NicolaDM/MAPLE). The logarithms of HnZ scores *H*(*n*) (see eqs. 1 and 2) are stored in a look-up table (nodes of the same size *n* have the same score) to reduce time demand. Node sizes *n* are also recorded and only updated when the size of a node is modified by a subtree prune and regraft[25] (SPR) move, a genome placement, or a branch length change. A threshold (equal by default to a tenth of the multiplicative inverse of the genome length) is also used, below which a branch length is considered effectively of length 0.

### 4.2 Usage of HnZ

MAPLE was always run with options “*--rateVariation --model UNREST*” specifying an UNREST[42] substitution model with rate variation[29].

For the analysis of real data, I used MAPLE v0.7.1 with HnZ1 (“*--HnZ 1*”). Phylogenetic inference was divided in four stages:

- Initial tree inference (“*--noFastTopologyInitialSearch --numTopologyImprovements 0*”) with one core (≈8.5 days and 20Gb).
- First topological improvement (“*--fast*”), using the tree from the previous step as initial tree (“*--largeUpdate --inputTree*”), and 14 cores (“*--numCores 14*”, ≈10 days and 500Gb).
- Further topological improvement (“*--noFastTopologyInitialSearch --thresholdLogLKtopology*
- *--allowedFailsTopology 8*”), using the tree from the previous step as initial tree (“*--largeUpdate --inputTree*”), and 14 cores (“*--numCores 14*”, ≈5.4 days and 450Gb).
- In-depth uncertainty assessment with SPRTA (“*--SPRTA --supportFor0Branches --numTopologyImprovements 0 --networkOutput --doNotImproveTopology --estimateMAT*”), using the tree from the previous step as initial tree (“*--largeUpdate --inputTree*”), and 1 core (≈12 days and 30Gb).

For simulated data, MAPLE v0.7.5.4 was run in a single step and on a single core either with HnZ (“*--HnZ 1*” or “*--HnZ 2*”) or without HnZ.

### 4.3 Datasets

#### 4.3.1 Real SARS-CoV-2 genome dataset

The real SARS-CoV-2 genome dataset considered in section 2.3 contains 2,072,111 genomes collected up to February 2023 and called with Viridian[43]. This dataset was collected and processed in [29]; in particular, potentially contaminated samples (those containing high level of heterozygosity) were filtered out, and alignment columns affected by recurrent sequence errors were masked. The alignment, metadata, inferred tree, and SPRTA support scores are available on Zenodo[44].

#### 4.3.2 Simulated genomes

For the benchmark of HnZ in Section 2.2 I used the SARS-CoV-2 genome data simulated in [29]. These genomes were evolved along a known (“true”) background phylogeny, the publicly available 26 October 2021 global SARS-CoV-2 phylogenetic tree from http://hgdownload.soe.ucsc.edu/goldenPath/wuhCor1/UShER_SARS-CoV-2/[45] representing the evolutionary relationship of 2,250,054 SARS-CoV-2 genomes, as inferred using UShER[22]. phastSim v0.0.3[46] was used to simulate sequence evolution along this tree according to SARS-CoV-2 non-stationary neutral mutation rates[28], using the SARS-CoV-2 Wuhan-Hu-1 genome[47] as root sequence, and with gamma-distributed (*α* = 0.2) rate variation[48], similar to that estimated from real data[29]. Ambiguity characters and sequence errors were then introduced in the alignment to mimic sequence incompleteness observed in real SARS-CoV-2 genome data and inferred recurrent errors. See [29] for a full description of these simulations.

## 5 Data Availability

The annotated SARS-CoV-2 phylogenetic tree inferred with HnZ1 is publicly available on Zenodo[44] version 6 https://doi.org/10.5281/zenodo.19002275, together with the considered SARS-CoV-2 genome alignment.

The MAPLE code implementing HnZ that was used for the results in this manuscript is open source and available from https://github.com/NicolaDM/MAPLE.

## 6 Acknowledgments

I thank Nick Goldman and Samuel Martin for helpful discussions regarding this work.

N.D.M. was supported by the European Molecular Biology Laboratory (EMBL) and by MRC grant MR/Z503769/1.

